# Topological Analysis of SARS CoV-2 Main Protease

**DOI:** 10.1101/2020.04.03.023887

**Authors:** Ernesto Estrada

## Abstract

There is an urgent necessity of effective medication against SARS CoV-2, which is producing the COVID-19 pandemic across the world. Its main protease (M^pro^) represents an attractive pharmacological target due to its involvement in essential viral functions. The crystal structure of free M^pro^ shows a large structural resemblance with the main protease of SARS CoV (nowadays known as SARS CoV-1). Here we report that as average SARS CoV-2 M^pro^ is 1900% more sensitive than SARS CoV-1 M^pro^ in transmitting tiny structural changes across the whole protein through long-range interactions. The largest sensitivity of M^pro^ to structural perturbations is located exactly around the catalytic site Cys-145, and coincides with the binding site of strong inhibitors. These findings, based on a simplified representation of the protein as a residue network, may help in designing potent inhibitors of SARS CoV-2 M^pro^.

The main protease of the new coronavirus SARS CoV-2 represents one of the most important targets for the antiviral pharmacological actions againsts COVID-19. This enzyme is essential for the virus due to its proteolytic processing of polyproteins. Here we discover that the main protease of SARS CoV-2 is topologically very similar to that of the SARS CoV-1. This is not surprising taking into account that both proteases differ only in 12 amino acids. However, we remarkable found a topological property of SARS CoV-2 that has increased in more than 1900% repect to its SARS CoV-1 analogue. This property reflects the capacity of the new protease of transmitting perturbations across its domains using long-range interactions. Also remarkable is the fact that the amino acids displaying such increased sensitivity to perturbations are around the binding site of the new protease, and close to its catalytic site. We also show that this sensititivy to perturbations is related to the effects of powerful protease inhibitors. In fact, the strongest inhibitors of the SARS CoV-2 main protease are those that produce the least change of this capacity of transmitting perturbations across the protein. We think that these findings may help in the design of new potent anti-SARS CoV-2 inhibitors.

## 1 Introduction

Since December 2019 an outbreak of pulmonary disease has been expanding from the city of Wuhan, Hubei province of China [1, 2]. This disease–produced by a new coronavirus named SARS-CoV-2 [3]–has become pandemic in about three months, affecting more than 200 countries around the world. SARS-CoV-2 belongs to the genus *Betacoronavirus* [4, 5], to which the virus which produced the respiratory epidemic of 2003 (nowadays known as SARS-CoV-1) also belongs to. The new coronavirus shares about 82% of its genome with SARS CoV-1. In spite of this similarity and of the fact that SARS-CoV-1 appeared almost 20 years ago, there are currently no approved specific drugs against SARS-CoV-2 [6, 7, 8, 9]. In consequence, most of the clinical treatment used against the disease is symptomatic in combination with some repurposed drugs, such as the antiviral Remdesivir or the antimalarials chloquine [10] and hidroxychloroquine [11]. This situation urges the scientific community to search for specific antiviral therapeutics and vaccines against SARS-CoV-2.

An attractive pharmacological target against the novel coronavirus is its viral protease, also known as the main protease (M^pro^) of SARS CoV-2. It is a key enzyme for the virus because it is essential for proteolytic processing of polyproteins [12]. As remarked by Zhang et al. [13] “inhibiting the activity of this enzyme would block viral replication. Since no human proteases with a similar cleavage specificity are known, inhibitors are unlikely to be toxic.” The three-dimensional structure of SARS CoV-2 M^pro^ has been resolved at different resolutions [14, 15, 13]. Other structures of SARS CoV-2 M^pro^ complexed with inhibitors have also been reported in recent works [13, 16, 17].

There are some remarkable characteristics of SARS CoV-2 M^pro^ in relation to the protease of SARS CoV-1. They share 96% of amino acids sequence, i.e., they differ in the amino acids at only 12 out of 303 positions in the sequence. Zhang et al. [13] have reported that the superposition of the chain A of two structures corresponding to the main proteases of SARS CoV-1 and of SARS CoV-2, namely 2BX4 and 6Y2E, respectively, shows a root mean square (r.m.s.) deviation of only 0.53 Å for all C_α_ positions. The first question that emerges here is whether such similarities are also reflected at the topological structural level of the proteins. By topological we mean here the discrete topology emerging from a network theoretic representation of a protein. In this representation of the protein structure the nodes of the network represent amino acids and the edges connecting them indicate that the corresponding residues are at a distance in which they can interact to each other. Because the Euclidean distance between the amino acids is used to construct the network we more correctly should refer to this framework as topographical more than topological. This network theoretic representation has been previously used to answer several questions related to protein structure and functioning [18, 19, 20, 21, 22]. Among the tools in use, the one of node centrality [23, 24] has played a fundamental role (see for instance [22]. These indices capture the relative importance–both structural and dynamical–of an individual amino acid in the protein.

Here we construct protein residue networks (PRN) for SARS CoV-2 M^pro^ and some of its inhibitors. The PRN of SARS CoV-2 M^pro^ is illustrated in Fig. 1. We then analyze the similarities in the topological structure of SARS CoV-2 M^pro^ with that of SARS CoV-1 for which we also construct the corresponding PRN. We then show that both proteases are very similar in relation to a few topological characteristics which account for a very close environment around the amino acids. That is, when the measures used account for the locality of the topological environment of a residue the two proteases do not differ in more than 2%. However, when the measures considered account for wider environments around the nodes the difference between the two proteins can increase up to 10-20%. These measures quantify how a perturbation at an amino acid is transmitted through the whole structure to the rest of the residues in the protein. When this transmission is allowed not only between close pairs of amino acids but also between very distant ones, the difference between the two proteases increases up to 1900%. That is, SARS CoV-2 M^pro^ is 1900% more sensitive to the transmission of perturbations between amino acids through the topological structure of the protein than SARS CoV-1 M^pro^. We discovered that the residues with this largest sensitivity in SARS CoV-2 M^pro^ are the ones involved in the binding of the three inhibitors studied here. That is, the most central amino acids according to this long-range indices are also the most affected by the interaction with the inhibitors as they are either in the binding site or very close to it. Consequently, we have discovered that the most relevant amino acids from the topological point of view are also the most relevant ones for the binding of some inhibitors to the SARS CoV-2 M^pro^ and should play an important role in the design of drugs inhibiting this protease.

**Figure 1:**
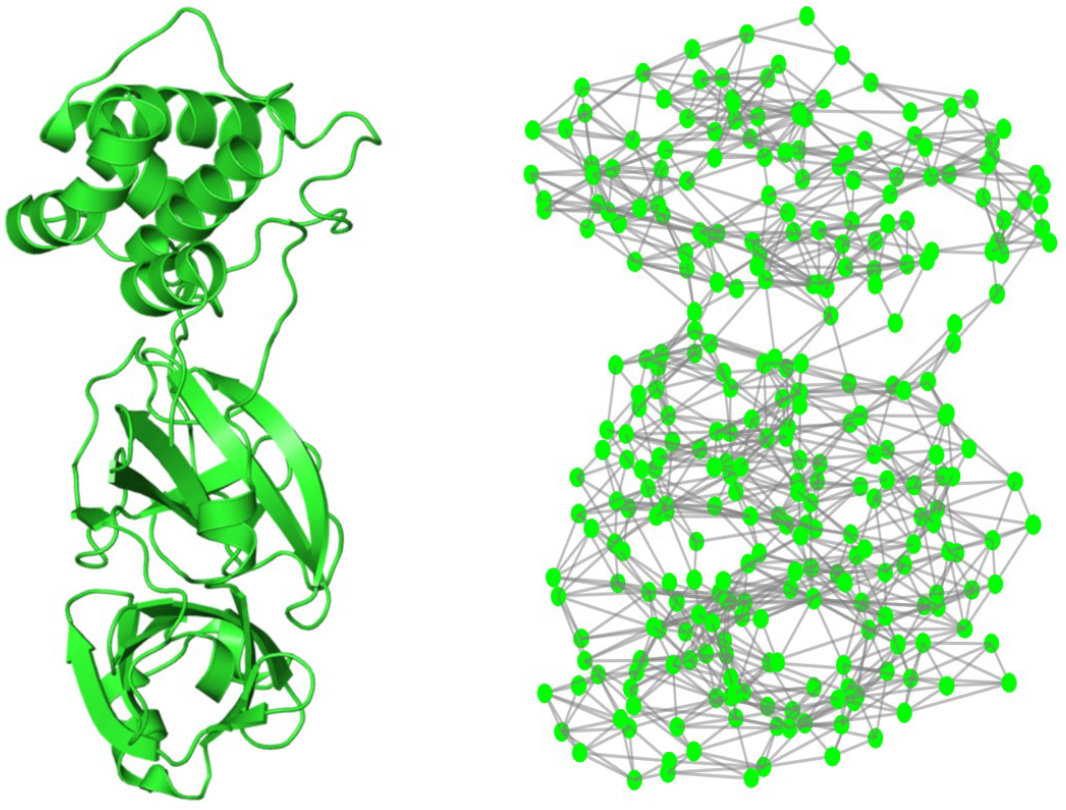
Cartoon representation (left) of the M^pro^ of SARS CoV-2 (PDB=6Y2E) and the corresponding protein residue network (right).

## 2 Methods

### 2.1 Construction of the protein residue networks

The protein residue networks (PRN) (see ref. [23] Chapter 14 for details) are built here by using the information reported on the Protein Data Bank [25] for the proteases of SARS CoV-1 and SARS CoV-2 as well as the complexes of the last one with an inhibitor. The nodes of the network represent the α-carbon of the amino acids. Then, we consider cutoff radius *r_C_*, which represents an upper limit for the separation between two residues in contact. The distance *r_ij_* between two residues *i* and *j* is measured by taking the distance between C_α_ atoms of both residues. Then, when the inter-residue distance is equal or less than *r_C_* both residues are considered to be interacting and they are connected in the PRN. The adjacency matrix *A* of the PRN is then built with elements defined by

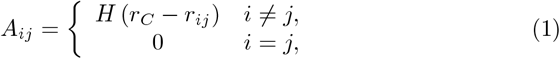

where *H*(*x*) is the Heaviside function. Here we use the typical interaction distance between two amino acids, which is equal to 7.0 Å. We have tested distances below and over this threshold obtaining in general networks which are either too sparse or too dense, respectively.

In this work we consider the structures of the M^pro^ of SARS CoV-1 deposited in the PDB with codes: 2H2Z [26], 2DUC [27], 1UJ1 [28], and 2BX4 [29]. We also study the following structures of of SARS CoV-2 with PDB codes: 6M03 [14], 6M2Q [14], and 6Y2E [13]. For the complexes of M^pro^ of SARS CoV-2 with inhibitors we study the structures with PDB codes: 6M0K [17], 6YZE [17] and 6Y2G [13].

The length of the proteases is 306 amino acids. However, there are structures (see Table 1) which are only resolved for amino acids 3 to 300, which gives a length of 298 [29]. Thus, for the sake of homogeneity of the analysis we consider here the same part of the amino acids sequence for all the structures analyzed, i.e., from residue 3 to residue 300. This does not alter the analysis as the two extremes of the protease are disordered and do not participate in important interactions.

**Table 1:** Protein Data Bank codes for structure of the main protease of SARS CoV-1 and SARS CoV-2 without inhibitors (apo forms). In the structure 1Q2W the residues 45-48 are missing. **The structure 3VB3 is resolved with two ligands di(hydroxyethyl)ether and 1,2-ethanediol. ***The structure 6YB7 is resolved with dimethylsulfoxide as a ligand.

| SARS CoV-1 |  |  | SARS CoV-2 |  |  |
| --- | --- | --- | --- | --- | --- |
| PDB | res. (Å) | length | PDB | res. (Å) | length |
| 2H2Z | 1.60 | 306 | 6YB7*** | 1.25 | 306 |
| 2DUC | 1.70 | 306 | 6M2Q | 1.70 | 305 |
| 1Q2W* | 1.86 | 295 | 6Y2E | 1.75 | 306 |
| 1UJ1 | 1.90 | 301 | 6M03 | 2.00 | 306 |
| 3VB3** | 2.20 | 301 |  |  |  |
| 2BX4 | 2.79 | 298 |  |  |  |

### 2.2 Network measures

The first category of measures correspond to those related to the most local structure around the nodes, such as those based on the degree of the nodes, i.e., the number of connections that a node has (see [23] for details). The degree accounts for the immediate effect of a node to its closest neighborhood. Among these measures we use here the edge density, which is defined as 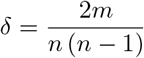, where *m* is the number of edges and *n* is the number of nodes. Because the average degree 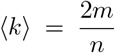, the relation with the edge density is clear. Another measure related to the degree of the nodes is the degree heterogeneity, 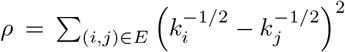 [30], which represents a measure of how heterogeneous the degrees of the nodes is [31]. A regular network, i.e., a network with all nodes of the same degree, will have *ρ* = 0, it is followed by networks with normal-like degree distributions, then networks with more heterogeneous ones, and will end up with networks with in which the probability *P*(*k*) of finding a node of degree *k* decays like distribution of the form *P*(*k*) ~ *k*^−1^, where *ρ* = 1. The average Watts-Strogatz clustering coefficient [32] is defined as 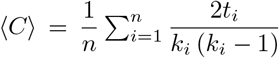, where *t_i_* is the number of triangles incident to the vertex. It account for the cliquishness around a node in terms of triangles, that is it account for how crowded the immediate neighborhood of a node is. We use Newman modularity index *Q* [33] to account for the modular structure of PRNs. It is defined as [33]: 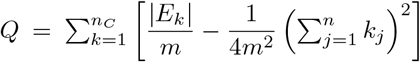, where |*E_k_*| is the number of edges between nodes in the *k*th community of the network, *m* is the total number of edges in the network and *k_j_* is the degree of the node *j*. these communities were previously detected by using Newman eigenvector method [34]. Another measure related to the degree is the degree assortativity coefficient [35], which is Pearson correlation coefficient of the degree-degree correlation. *r* > 0 (degree assortativity) indicates a tendency of high degree nodes to connect to other high degree ones. *r* < 0 (degree disassortativity) indicates the tendency of high degree nodes to be connected to low degree ones. Other measures in this class assume that “information” is transmitted in the network through the topological shortest paths. The length of the shortest path is a distance *d*(*i,j*) between the corresponding pairs of nodes *i* and *j*, and it is known as the shortest path distance. The average path length 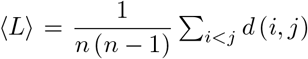 is typically used as a measure of the ‘small-worldness’ of the network [32]. We also consider the average betweenness centrality [36] 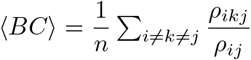, where *ρ_ikj_* is the number of shortest paths between the nodes *i* and *j* that cross the node *k*, and *ρ_ij_* is the total number of shortest paths that go from *i* to *j*. It accounts for the importance of a node in passing information through it to connect other pairs of nodes via shortest path only.

The second category of measures is formed by those that account for the transmission of information not only via the shortest paths but by using any available route that connect the corresponding pair of nodes. These measures use the concept of walk instead of that of a path. A walk of length *k* in *G* is a set of nodes *i*_1_, *i*_2_,…, *i_k_*, *i*_*k*+1_ such that for all 1 ≤ *l* ≤ *k*, (*i_l_*, *i*_*l*+1_) ∈ *E*. A *closed walk* is a walk for which *i*_1_ = *i*_*k*+1_. The number of walks of length *k* between the nodes *i* and *j* in a network is given by (*A^k^*)_*ij*_. The first of these measures considered here is the eigenvector centrality *EC* [37], which is the corresponding entry of the eigenvector associated with the largest eigenvalue of *A*. The relation of this index with walks is given by the following. Let *N_k_*(*i*) be the number of walks of length *k* starting at node *i* and ending elsewhere. Then, if the network is not bipartite, which is the case of the current work, 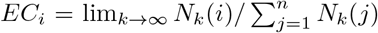 (see Chapter 5 in [23]). That is, the eigenvector centrality of a node is the ratio of the number of walks of infinite length that start at this node to the whole number of such walks starting elsewhere. Consequently, the average eigenvector centrality 〈*EC*〉, accounts for the spread of information from the nodes beyond the nearest neighbors and using any infinite-length walk in the graph. A type of measures of the second kind are based on counting all walks of any length, but giving more weight to the shorter than to the longer ones. These measures are based on the following matrix function: 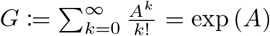, where exp(*A*) is the exponential of the matrix. then, we consider the average of the diagonal entries of this matrix, which is known as the average subgraph centrality 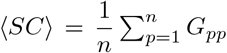 [38], which accounts for the participation of the corresponding node in all subgraphs of the graphs, giving more weight to the shortest than to the longer ones. Such subgraphs include for instance, edges, triangles, wedges, squares, etc. Another measure is the average of the non-diagonal entries of exp(*A*), which is known as 2 the average communicability of the network, 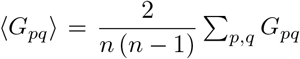[39].

It accounts for how much a pair of nodes can communicate to each other by using all potential routes available in the network, but giving more weight to the shortest than to the longer ones. Finally, in this category we include the average communicability angle 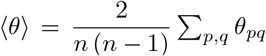 [40], where the angle between a pair of nodes is defined as: 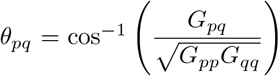. The average communicability angle describes how efficiently a network transmit information between its pairs of nodes by using all available routes.

The third category of measures is formed by all-walks indices that penalize less heavily longer walks connecting pairs of nodes in a network. That is, although *G* = exp(*A*) accounts for all walks connecting every pair of nodes, it penalizes very much those walks of relatively large length, then making more emphasis in shorter walks around a given node. In order to include longer walks in the analysis we study the following matrix function [41]: 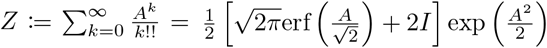, which penalizes the walks of length *k* not by *k*! (simple factorial) but by *k*!! (double factorial). Then, we will consider here the average of the main diagonal 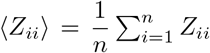, which accounts for the participation of the node *i* in all subgraphs in the graph but including bigger subgraphs than in *SC*. In a similar way we consider 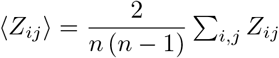, which accounts for the global capacity of the network of transmitting information between pairs of nodes and allowing longer-range transmission than in the case of the communicability. For those reasons we propose to call these indices long-range (LR) subgraph centrality and communicability.

## 3 Results

### 3.1 Free protease

The main goal of this section is to analyze a few network theoretic measures of the M^pro^ of SARS CoV-2 and compare them with those of the protease of SARS SARS CoV-1. The amino acid sequence of both proteases share 96% of similarity, i.e., only 12 amino acids are different in both proteases of a total of 303. These amino acids are at positions 33, 44, 63, 84 86, 92, 132, 178, 200, 265, 283 and 284. In order to compare the topological features of the main proteases of SARS CoV-1 and of SARS CoV-2 we go a step further here and compare several structures of the M^pro^ of SARS CoV-1 and SARS CoV-2. In Table 1 we give the PDB codes of 6 structures of the main protease of SARS CoV-1 and 4 of SARS CoV-2 without inhibitors. In these structures not only there are no inhibitors, but also there are no mutations in the structure of the wild proteases. In the case of the structure with PDB code 1Q2W the residues 45-48 are missing in the PDB. In 3VB3 we have found that di(hydroxyethyl)ether (PEG) and 1,2-ethanediol (EDO) are also present in the crystal structure. In a similar way the structure 6YB7 contains dimethylsulfoxide (DMS) in the crystal structure. For these reasons we will not include these three structures in the further analysis.

For the rest of the structures, i.e., 4 structures of the main protease of SARS CoV-1 and 3 structures of the same for SARS CoV-2, we calculate all the topological measures defined in Methods. We then obtained the mean and standard deviation of these measures for the two groups of structures and report them in Table 2. We can observe in this Table that most of the topological characteristics of the first kind of the PRNs of both proteases are very similar with relative differences not bigger than 2% for all the properties analyzed. In order to test the significance of the differences between the two groups of proteases we use the p-values of the Mann-Whitney U-test [42]. This statistical measure has been proposed for the analysis of network measures, in particular for protein networks [43, 44]. According to the p-values (see last column in Table 2) none of these measures display significant difference between the two groups of proteases.

**Table 2:** Average values of the global topological properties of the M^pro^ of SARS CoV-1 (2H2Z, 1UJ1, 2DUC, 2BX4) and SARS CoV-2 (6M03, 6Y2E, 6M2Q). The relative difference between them, expressed as percentages of change relative to SARS CoV-1, and the p-values of the Mann-Whitney U test are also given.

| measure | SARS CoV-1 | SARS CoV-2 | $\Delta_{rel}$ (%) | U-stat |
| --- | --- | --- | --- | --- |
| $\delta$ | 0.0260 | 0.0262 | -0.71 | 0.2286 |
| $\rho$ | 0.0163 | 0.0164 | -0.98 | 0.8571 |
| $\langle L \rangle$ | 6.37 | 6.33 | 0.69 | 0.2286 |
| $Q$ | 0.613 | 0.610 | 0.49 | 1.0000 |
| $\langle C \rangle$ | 0.542 | 0.540 | 0.31 | 0.8571 |
| $r$ | 0.390 | 0.398 | -1.85 | 0.8571 |
| $\langle BC \rangle$ | 796.29 | 793.76 | 0.32 | 0.6286 |
| $\langle EC \rangle$ | 0.00336 | 0.00334 | 0.50 | 0.5714 |
| $\langle SC \rangle$ | 172.00 | 196.04 | **-13.97** | **0.0571** |
| $\langle G_{pq} \rangle$ | 22.42 | 26.46 | **-18.01** | **0.0571** |
| $\langle \theta \rangle$ | 82.29 | 82.01 | 0.34 | **0.0571** |
| $\langle Z_{pp} \rangle$ | $4.65 \cdot 10^{17}$ | $9.57 \cdot 10^{18}$ | **-1960.15** | **0.0571** |
| $\langle Z_{pq} \rangle$ | $1.44 \cdot 10^{17}$ | $2.91 \cdot 10^{18}$ | **-1921.88** | **0.0571** |

We then continue the analysis by comparing the topological measures of the second kind. We notice that the eigenvector centrality, which has been found very useful in previous analysis of PRN [22], does not display any significant difference between both proteases according to the Mann-Whitney test. However, there are differences in the mean subgraph centrality of about 14% and of the average communicability between pairs of nodes of about 18%. In both cases, the indices are significantly larger for the protease of SARS CoV-2 than for that of SARS CoV-1. According to the p-values these differences are significant at 94% level of confidence in the Mann-Whitney U-test. This means that the structural changes that make the difference between the proteases of SARS CoV-1 and SARS CoV-2 increase the capacity of the individual amino acids of feeling a perturbation or thermal oscillation produced in another amino acid of the protein. As we have previously explained these communicability factors penalizes very heavily any perturbation being transmitted between two amino acids separated by a relatively long distance in the protein. Thus, they can be considered as indices that account for shorter range interactions than the third kind measures considered here. It should be noticed that although the communicability angles display very little relative variation between the two groups of proteases, these differences are significant at 94% of confidence in the Mann-Whitney test.

Both LR subgraph centrality and communicability display dramatic increment in SARS CoV-2 relative to SARS CoV-1. In this case the increase of these indices is more than 1900% for both, the LR communicability and LR subgraph centrality. In short, this means that the protease of SARS CoV-2 has more than 13 times more capacity of transmitting perturbations between pairs of nodes than the protease of SARS CoV-1. This is equivalent to say that the protease of SARS CoV-2 is significantly much more topologically efficient in transmitting “information” among its amino acids than the protease of SARS CoV-1. These two topological measures display significant differences between the two groups of proteases according to the statistical p-values obtained from the Mann-Whitney U-test at 94% of confidence.

We now proceed to the analysis of the local variation of the subgraph and the LR subgraph centralities for the amino acids of the two M^pro^ (see Fig. 2) averaged for all the structures previously mentioned, i.e., 2H2Z, 1UJ1, 2DUC, 2BX4 for SARS CoV-1 and 6M03, 6Y2E, 6M2Q for SARS CoV-2. In the case of the subgraph centrality the largest change is produced for a few amino acids which increase their centrality in SARS CoV-2 relative to SARS CoV-1. These are the cases of 25, 26, 27, 118, 17, and 24. But there are also other amino acids which drop their centrality in SARS CoV-2, such as 170, 73, 169, 165, 89, and 252 among others (see Fig. 2(b)). Therefore, the increase of the subgraph centrality of a few amino acids makes that in total the average subgraph centrality increases in SARS CoV-2 in relation to SARS CoV-1. An important characteristic feature of the differences in this centrality between the two proteases is that they are spread across the three domains of the proteases with a large increment in the domains I and III. This is a major difference with the LR subgraph centrality (see Figs. 2(c) and (d)), where the main change is a dramatic increase in the centrality of the nodes in the domains I and II of the SARS CoV-2 protease relative to SARS CoV-1. The changes occurring in the domain III are imperceptible in relation to those of the other two domains.32

**Figure 2:**
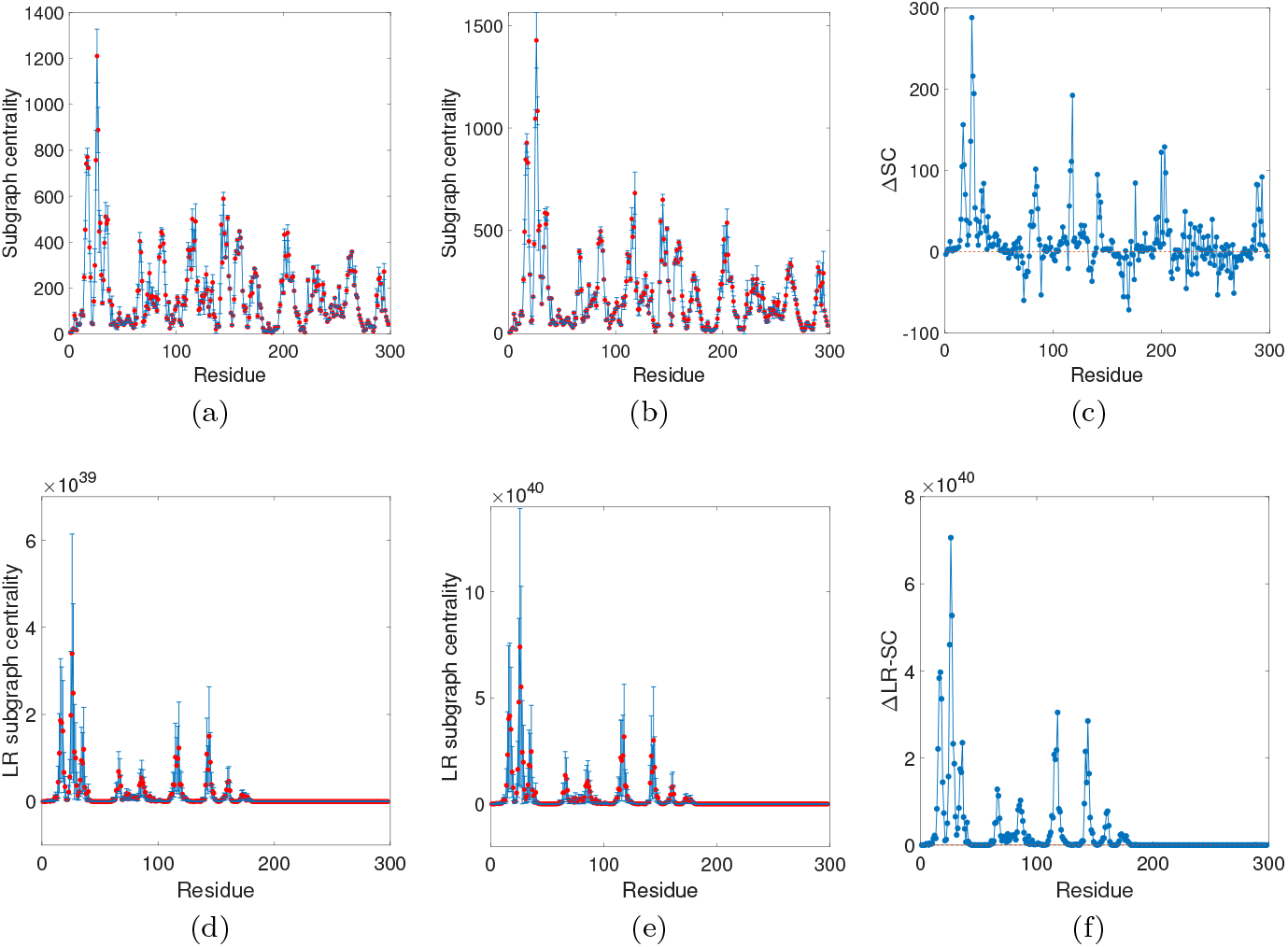
Plot of topological properties of the amino acid residues for the M^pro^ of SARS CoV-1 (broken red line) and of SARS CoV-2 (solid blue lines). (a) Subgraph centrality; (b) LR subgraph centrality.

In order to illustrate the distributions of the most central amino acids according to both measures in the three-dimensional structures of the proteases we selected two structures, 2BX4 for SARS CoV-1 and 6Y2E for SARS CoV-2 as representative of the two groups of structures. Notice that these two structures have been used by Zhang et al. [13] for their comparison of the 3D structures of both proteases. Both structures are illustrated in Fig. 3. It can be seen that the largest values of the LR subgraph centrality are concentrated in a relatively small region of the protein structure, while those of the subgraph centrality are more spread across the whole structure. We then inquire about this region of the M^pro^ in SARS CoV-2 which shows the largest change in the LR subgraph centrality relative to its analogue of SARS CoV-1.

**Figure 3:**
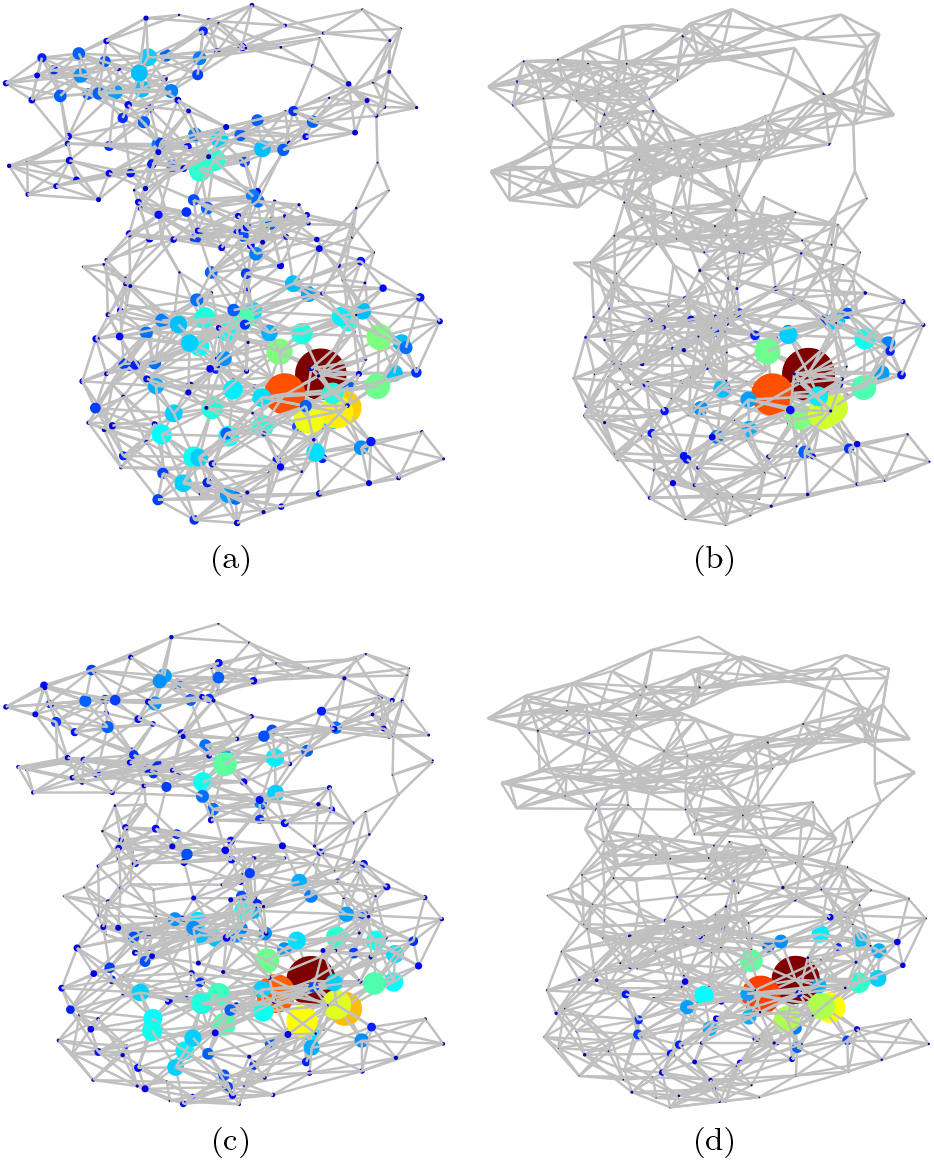
Illustration of the subgraph (a), (c) and LR subgraph (b), (d) centralities of the amino acid residues of the chain A of SARS CoV-1 M^pro^ of (top), and of SARS CoV-2 (bottom). The size of the nodes is proportional to the corresponding centrality normalized to its largest value in the protease analyzed. The colors also correspond to the same values in the jet color code, with red for higher and blue for smaller values.

The first remarkable observation of the amino acids with the largest change in the LR subgraph centrality is that they are all closely separated to each other in the three-dimensional space. For instance, the 22 amino acids displaying the largest change in this centrality form a connected subgraph of the PRN as illustrated in Fig. 4. This subgraph of 22 nodes has 48 connections among these amino acids, which produces an edge density of 0.21, almost 10 times bigger than the total density of the protease. The second remarkable feature of this subgraph is that it contains one of the two catalytic amino acids of the M^pro^ of SARS CoV-2, which is Cys-145. That is, the region with the largest increase in the LR subgraph centrality of the protease of SARS CoV-2 relative to SARS CoV-1 is the one enclosing the catalytic binding site of amino acid Cys-145. It is also remarkable that this region of large increment in the LR subgraph centrality contains some amino acids which are located in the binding site of the M^pro^ to α-ketoamide inhibitors as well as other kind of inhibitors, as we will analyze further in this work. This is the case of the residue 144-147, other amino acids in this binding site like residues 162, 163 also display large increment in the LR subgraph centrality. The last remarkable observation is that the domain III displays small change in relation to the changes of domains I and II in this topological parameter. However, as we will see in the next paragraphs this domain (residues 198-303) which is formed by 5 helices and is involved in the dimerization of the M^pro^, also increases significantly the LR communicability in relation to SARS CoV-1.

**Figure 4:**
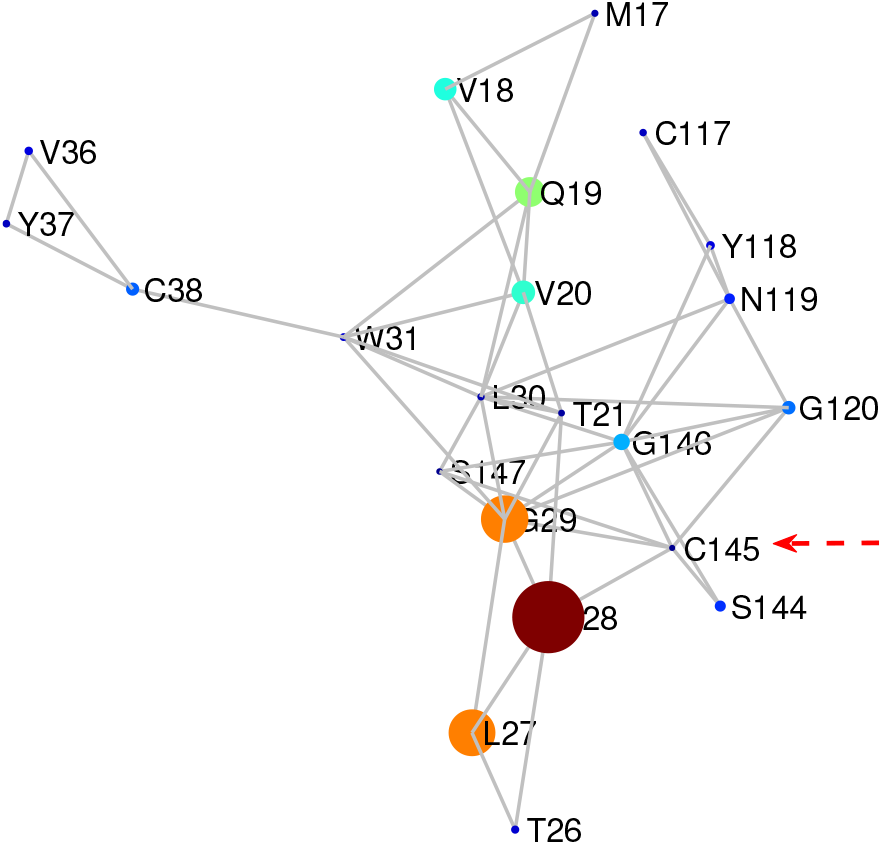
Illustration of the 22 amino acids which display the largest difference in the LR subgraph centrality in a representative M^pro^ of SARS CoV-2 (6Y2E) in relation to one of SARS CoV-1 (2BX4). The radius of the nodes is proportional to the difference in the LR subgraph centrality between the two proteases. The catalytic site Cys-145 is pointed to with an arrow. The size of the nodes is proportional to the corresponding centrality normalized to its largest value in the protease analyzed. The colors also correspond to the same values in the jet color code, with red for higher and blue for smaller values.

A better picture of the changes in the different regions of the M^pro^ of SARSCoV-2 relative to SARS CoV-1 can be obtained again by analyzing the differences between the communicabilities and LR-communicabilities averaged for the 4 structures of SARS CoV-1 and 3 structures of SARS CoV-2 before considered. For this we obtain an average communicability (resp. LR communicability) matrix for the structures of SARS CoV-1 and another for the structures of SARS CoV-2. Then, we obtain the difference between these two matrices. In Fig. 5 we illustrate the difference matrices for both kinds of communicabilities. In the first case it can be observed that the communicability between all pairs of residues in the domain I (residues 10-99) mainly increase in SARS CoV-2 relative to SARS CoV-1, with an increase of 12.8% relative to SARS CoV-1. However, in the domain II (residues 100-182) there is mainly a drop of the communicability between the residues in the domain, which decrease 2.02%, but there is an increase of 19.6% in the trade off between domains I and II, and an increase of 39.2% in the trade between domains I and III. The domain III shows a mixed behavior with some pairs of residues increasing and other decreasing their communicability, but the main result is an increase of 5.58% relative to SARS CoV-1. The communicability between domains II and III in the SARS CoV-2 structures increase in 23.9% relative to the same in SARS CoV-1.

**Figure 5:**
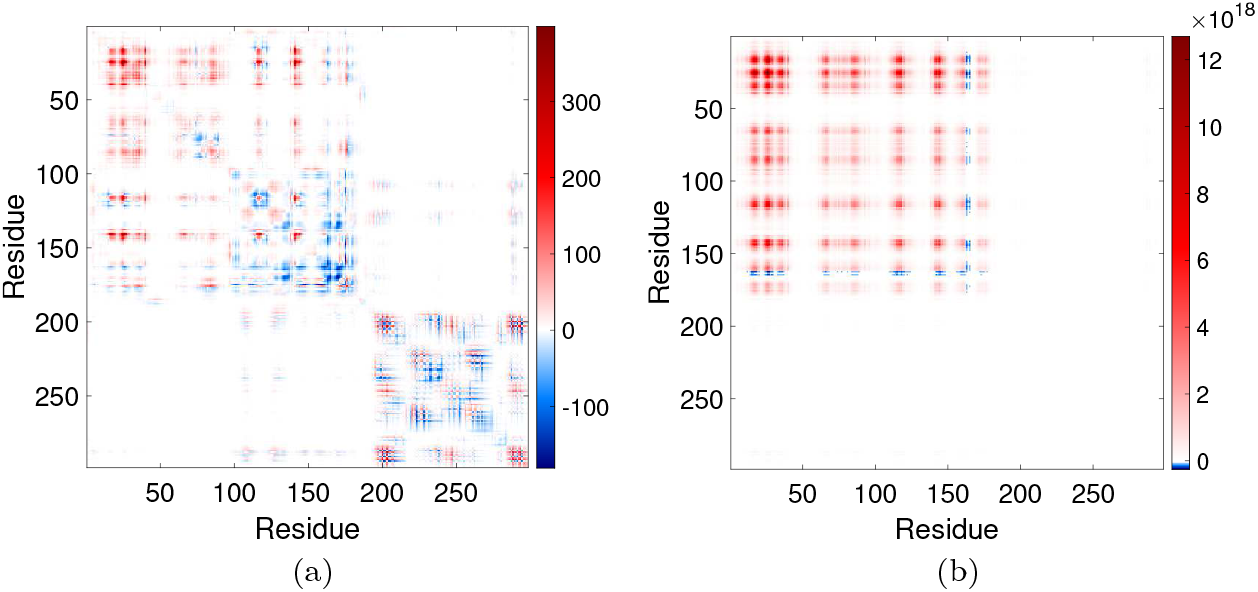
Difference between communicabilities (a), and LR communicability (b) between pairs of amino acids in the averaged structures of M^pro^ of SARS CoV-2 in relation to that of SARS CoV-1.

We finally analyze the changes in the LR communicability between the different domains of the SARS CoV-2 protease. Here the changes are dramatic and in all cases the LR communicability in the SARS CoV-2 protease is higher than that in SARS CoV-1. For instance, the average communicability between pairs of nodes in the domain I is 1997% higher in SARS CoV-2 than in SARS CoV-1. This percentage of increment are 1814% in the domain II and 2651% in domain III. The inter-domain communicability also increases very significantly with increment of 1896% (domains I-II), 2350% (domains (I-III) and 2237% (domains II-III). In closing, the structural changes between the main proteases of SARS CoV-1 and SARS CoV-2 produced a dramatic impact in the LR communicability between residues in the protease of SARS CoV-2 with huge improvement in long-range communication between residues practically in all domains of the protease.

### 3.2 SARS CoV-2 protease bounded to inhibitors

We turn now our attention to the analysis of the M^pro^ of SARS CoV-2 complexed with some inhibitors. The selection of these inhibitors has been based on: (i) the existence of the crystallographic structure of the complex inhibitor-M^pro^, (ii) the existence of reports about the inhibitory concentration *IC*_50_ of the inhibitor, and (iii) the fact that the inhibitors display a great potency against the main protease of SARS CoV-2. Then, we have selected three complexes which correspond to PDB codes 6M0K, 6LZE and 6Y2G. The first two compounds were recently reported by Dai et al. [17] and the third is an α-ketoamide inhibitor reported by Zhang et al. [13]. The first two inhibitors display *IC*_50_ < 0.1*μM* and the third shows *IC*_50_ ≈ 0.67 ± 0.18.

In Table 3 we resume the results of the calculation of average topological properties of the M^pro^ structure bounded to these inhibitors. In these calculations we consider only the residues 3-298 of the protease as explained in Methods to make these results comparable with the ones obtained in the previous section. It can be seen that here again the topological measures of the first class display relatively little variation for the three complexed proteases relative to the free one.

**Table 3:** Relative differences in percentage of global topological properties of the M^pro^ of SARS CoV-2 complexed to an inhibitor in relation to free one. The PDB of the complexes between the M^pro^ of SARS CoV-2 with an inhibitor correspond to 6Y2F (space group *C*_2_), and 6Y2G (space group *P*2_1_2_1_2_1_).

| measure |  | wild | 6M0K | 6LZE | 6Y2G |
| --- | --- | --- | --- | --- | --- |
| $\delta$ | | 0.0262 | 0.0262 | 0.0262 | 0.0255 |
| $\rho$ | | 0.0164 | 0.0167 | 0.0164 | 0.0178 |
| $\langle L \rangle$ | | 6.33 | 6.386 | 6.383 | 6.35 |
| $\langle C \rangle$ | | 0.54 | 0.54 | 0.54 | 0.54 |
| $r$ | | 0.398 | 0.394 | 0.375 | 0.37 |
| $\langle BC \rangle$ | | 793.76 | 799.86 | 799.39 | 795.92 |
| $\langle EC \rangle$ | | 0.00334 | 0.00335 | 0.00336 | 0.0033 |
| $\langle SC \rangle$ | | 196.04 | 187.85 | 180.42 | 156.09 |
| $\langle G_{pq} \rangle$ | | 26.46 | 25.07 | 23.40 | 20.09 |
| $\langle \theta \rangle$ | | 82.01 | 82.12 | 82.24 | 82.45 |
| $\langle Z_{ii} \rangle$ | | $9.57 \cdot 10^{18}$ | $1.99 \cdot 10^{18}$ | $4.79 \cdot 10^{17}$ | $1.28 \cdot 10^{17}$ |
| $\langle Z_{ij} \rangle$ | | $2.91 \cdot 10^{18}$ | $5.93 \cdot 10^{17}$ | $1.54 \cdot 10^{17}$ | $4.06 \cdot 10^{16}$ |
| $IC_{50} (\mu M)$ | | | $0.04 \pm 0.002$ | $0.053 \pm 0.005$ | $0.67 \pm 0.18$ |

We then move to the analysis of the measures of second and third type. As can be seen in Table 3 there are significant changes, of more than 20%, in the subgraph centrality and the communicability of the complexed proteases in relation to the average of the wild proteases previously analyzed. However, here again, the most dramatic change in these topological properties occurs in the values of the LR subgraph centrality and communicability, with relative changes of more than 98%. We should notice that the smallest change in these parameters occurs for the structure 6M0K, which corresponds to the strongest inhibitor, followed by 6LZE, which is the intermediate one, and finally 6Y2G which is the weakest of the three. That is, the strongest inhibitor produces the smallest changes in the (LR) subgraph centrality and (LR) communicability in relation to the wild protease. In contrast, the weakest inhibitor changes the most these communicability parameters relative to the unbounded protease. These results appears to indicate that the potency of these inhibitors could be related to the fact of not affecting very much the strong inter-residue communicability of amino acids in the M^pro^ of SARS CoV-2.

With the goal of disentangling the information contained in the changes produced at the LR subgraph centrality of the bounded protease we study it in more detail here. For this, we consider the amino acids displaying the largest values of this topological parameter for the three structures. In Fig. 6 we illustrate the region formed by the top 22 amino acids according to their values of the *Z_ij_* index, i.e., LR subgraph centrality. The first interesting observation is that for the three structures considered these amino acids form a connected subgraph in the main protease. That is, these amino acids displaying the highest LR subgraph centrality are not randomly distributed around the domains of the protease but they are located in a specific location of the space. It is also remarkable that this subgraph is connected, which means that there is no single amino acid separated at more than 7 Å from all the rest of residues forming the subgraph. Another remarkable characteristic of these subgraphs of the most central residues according to LR subgraph centrality is that they are exactly around the binding site of the main protease. As can be seen in the Fig. 6 these subgraphs of residues are very close to the inhibitors and form a cluster of amino acids around the catalytic site, which is C145.

**Figure 6:**
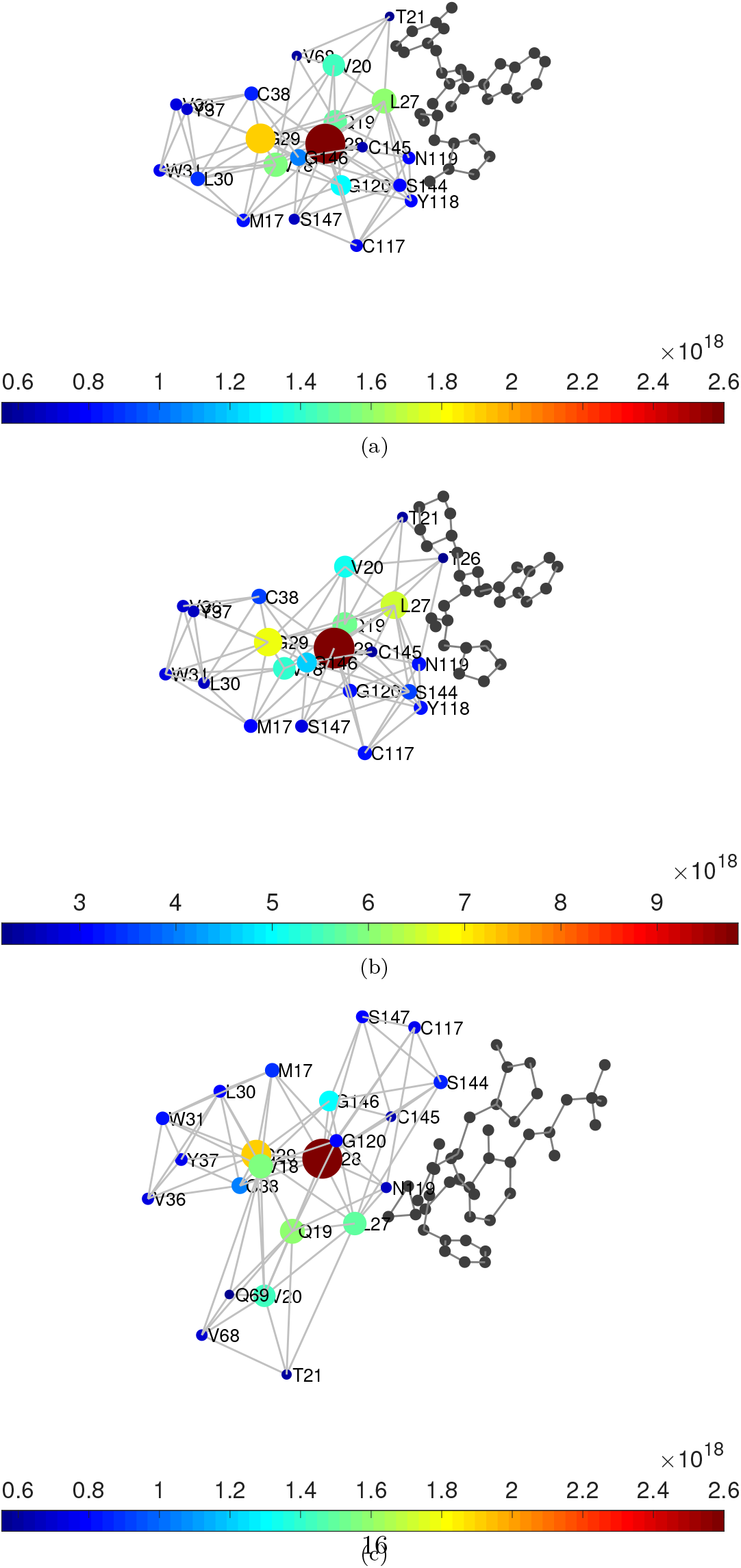
Illustration of the 22 amino acids with the largest values of the LR subgraph centrality in 6M0K (a), 6LZE (b) and 6Y2G (c). The residues are connected if they are at no more than 7.0 Å. The color bar and the radius of the nodes indicates the values of *Z_ii_* normalized to the largest value in the corresponding protein.

In Table 4 we resume the results of the top ranked amino acids according to the LR subgraph centrality for the free SARS CoV-2 M^pro^ taken as the average of the three apo structures previously considered and the three complexes with inhibitors studied here. As can be seen the top 22 amino acids in the average free SARS CoV-2 M^pro^ contains more than 90% of the residues which appear involved in the interactions with the three inhibitors studied here. In the case of 6LZE they coincide in 100%, and in 6M0K the coincidence is of 95%.

**Table 4:** List of amino acids with the largest values of LR subgraph centrality in the average free protease (average of 6M03, 6Y2E, 6M2Q) and with the same parameter for the protease bounded to inhibitors (6M0K, 6LZE, and 6Y2G). The amino acids in the bounded protease which are not in the top rank of the free one are marked in red.

| Rank | average | 6M0K | 6LZE | 6Y2G |
| --- | --- | --- | --- | --- |
| 1 | N28 | N28 | N28 | N28 |
| 2 | G29 | G29 | G29 | G29 |
| 3 | L27 | L27 | L27 | Q19 |
| 4 | V18 | V18 | Q19 | V18 |
| 5 | Q19 | Q19 | V18 | L27 |
| 6 | V20 | V20 | V20 | V20 |
| 7 | G146 | G120 | G146 | G146 |
| 8 | G120 | G146 | S144 | C38 |
| 9 | L30 | L30 | C38 | M17 |
| 10 | C38 | C38 | N119 | S144 |
| 11 | M17 | M17 | C117 | W31 |
| 12 | N119 | S144 | G120 | G120 |
| 13 | S144 | N119 | Y118 | S147 |
| 14 | Y118 | Y118 | M17 | L30 |
| 15 | C117 | C117 | S147 | Y37 |
| 16 | W31 | W31 | W31 | C117 |
| 17 | V36 | V36 | V36 | V36 |
| 18 | Y37 | Y37 | L30 | V68 |
| 19 | S147 | S147 | Y37 | N119 |
| 20 | T26 | C145 | T21 | C145 |
| 21 | C145 | V68 | C145 | T21 |
| 22 | T21 | T21 | T26 | Q69 |

## 4 Discussion and Conclusions

We present an analysis of some of the most relevant topological properties of the main protease of the SARS CoV-2. Our approach is based on the representation of the three-dimensional structure of the protein as a residue network in which *C_α_* of every amino acid is represented by a node of the network and two nodes are connected if the corresponding *C_α_* are at no more than 7.0 Å. We find here that the difference between most of the topological properties of the PRNs representing both proteases differ in less than 5%. If we exclude from the analysis the LR measures, then 70% of the topological measures shows only a small variation between the two proteases taking as the average of the properties of several structures representing each of the two proteases. In this situation it is certainly remarkable that there are topological measures which change in more than 1000% from one protease to the other. These are the cases of the LR subgraph centrality of the amino acids and of the LR communicability between pairs of them. The increase of these parameters in more than 1900% for SARS CoV-2 M^pro^ relative to SARS CoV-1 M^pro^ means that the structural changes that differentiate both proteases have created a huge increment in the efficiency of SARS CoV-2 M^pro^ in transmitting perturbations of any kind between the amino acids of the protein using all available routes of connection and allowing for long-distance transmission. To make clearer what this sensitivity means we are going to use a simple example. Let us consider a tiny perturbation on the structure of the proteases which prevent the interaction between the amino acids P9 and G11, which have been selected at random. In SARS CoV-1 M^pro^ (taking 2BX4 as an example) these amino acids are at 5.69 Å and in SARS CoV-2 M^pro^ (taking 6Y2E as an example) they are 6.48 Å apart. Thus, in both cases they are connected in the corresponding PRN. Let us consider that the perturbation remove this edge from the PRN of both proteases. The relative decrement of the average path length in SARS CoV-2 M^pro^ relative to SARS CoV-1 M^pro^ is almost imperceptible, i.e., 5.7%. In the case of the subgraph centrality it is of the same order, i.e., 3.4%. This means that according to these parameters SARS CoV-2 M^pro^ is as sensitive as SARS CoV-1 M^pro^ to perceive a structural change in its structure produced by a given perturbation. However, when we consider the LR subgraph centrality this relative change is 316.8%. That is, according to this topological parameter which takes into account long-range interactions, SARS CoV-2 M^pro^ is more than three times more sensitive to a tiny structural change than SARS CoV-1 M^pro^. This remarkable finding indicates that the 12 mutations produced in SARS CoV-1 M^pro^ makes the resulting SARS CoV-2 M^pro^ much more efficient in transmitting “information” through the protein skeleton using short and long-range routes.is proportional to the absolute value of this difference.

The second remarkable finding of the current work is that the largest changes in the LR subgraph centrality occurring in SARS CoV-2 M^pro^ relative to SARS CoV-1 M^pro^ do not spread equally across the whole structure of the protease. Instead, they are concentrated around a geometrical region which includes most of the amino acids involved in the binding site of the protease to inhibitors or close to it. One of the amino acids which has increased more dramatically its sensitivity to long-range transmission of information in SARS CoV-2 M^pro^ is Cys-145, which is one of the two catalytic sites of the protease, and the one involved in interactions with the inhibitors, such as the ones analyzed here. We have analyzed here three different inhibitors of SARS CoV-2 M^pro^ displaying very potent inhibitory capacity over the protease. In the three cases we have observed a significant variation in the LR subgraph centrality of the amino acids which were previously observed to have increased their LR sensitivity in the free protease. Therefore, these amino acids corresponds to those involved in the binding of these three inhibitors, showing that their increased topological role in the SARS CoV-2 M^pro^ also may play an important functional role in it.

The analysis of PRN is easier than the study of the whole protein structure. In this sense the PRN represents a simplified model of the three-dimensional structure of the protein. Typically, such simplification in the complexity of the representation of systems convey a loss in the structural information which is represented by the global system. In this case, however, we have shown that the use of a network representation of the proteins reveals some hidden patterns in their structure that were escaping to the analysis by using the global structure. To detect such important structural factors it is necessary to account for long-range interactions among the amino acids of the proteases, which are the ones revealing the their most important characteristics in terms of their sensitivity to tiny structural changes produced by local or global perturbations to the system. Such LR interactions revealed here the main differences between the proteases of SARS CoV-1 and SARS CoV-2, as well as the most important amino acids for the interaction with inhibitors, which may produce therapeutic candidates against COVID-19.

## Supporting information

Adjacency list of PRN

x,y,z coordinates of nodes in the PRN

## Data availability statement

The data that supports the findings of this study are available within the article [and its supplementary material].

